# Surveillance-embedded genomic outbreak resolution of methicillin-susceptible *Staphylococcus aureus* in a neonatal intensive care unit

**DOI:** 10.1101/584359

**Authors:** AJH Cremers, JPM Coolen, CP Bleeker-Rovers, ADJ van der Geest-Blankert, D Haverkate, H Hendriks, SSV Henriet, MA Huynen, E Kolwijck, D Liem, WJG Melchers, JW Rossen, J Zoll, A van Heijst, J Hopman, HFL Wertheim

## Abstract

**Background:** We observed an increase in methicillin-susceptible *Staphylococcus aureus* (MSSA) infections among neonates at a Dutch third level neonatal intensive care unit. Weekly surveillance data of MSSA carriage among neonates and cross-sectional screenings of health care workers (HCWs) were available for outbreak tracing. While traditional typing of MSSA isolates by staphylococcal protein A gene (*spa* typing) and Multiple-Locus Variable number tandem repeat Analysis (MLVA) suggested that nosocomial transmission had contributed to the infections, here they lacked the resolution to draw solid conclusions.

**Methods:** MSSA isolates from neonatal infections, carriage surveillance, and HCWs were subjected to whole-genome sequencing and compared by a series of automated tools including *de novo* assembly, identification and localization of high-quality single nucleotide polymorphisms, and in-depth analysis of subsets of isolates. Outbreaks were defined as isolates that were more closely related than was to be expected from the genetic diversity in background surveillance.

**Results:** Genomic analysis identified isolates that had been unjustly assigned to clusters based on MLVA typing, while *spa* typing was concordant but of insufficient resolution. Detailing particular subsets of isolates further improved resolution and although it provided evidence that HCWs were involved in multiple outbreaks, it alleviated heavy concerns about one particular HCW. Genomic clustering of isolates based on deviations from background surveillance matched epidemiological patient linkage. Compared to MLVA typing, the genomic analysis demonstrated more, shorter, and re-assorted nosocomial transmission chains during this outbreak.

**Conclusions:** In this study the improved resolution and accuracy of genomic outbreak analyses compared to *spa* typing and MLVA substantially altered the view on outbreaks, along with apposite outbreak measures. Inclusion of the circulating background population has the potential to overcome current issues in genomic outbreak inference.

## Background

In the third level neonatal intensive care unit (NICU) of our hospital, we observed a rise in methicillin-susceptible *S. aureus* (MSSA) infections in two consecutive years. Traditional MSSA typing methods brought the suspicion of health care workers (HCWs) being repeatedly involved in the outbreak transmission chains. However, as these traditional typing methods fell short in providing definitive answers, we set out to improve the resolution of our outbreak analysis by bacterial whole-genome sequencing (WGS).

Neonates have an immature immune system and are susceptible to opportunistic bacterial infections, especially in case of premature birth. Important bacteria involved in outbreaks at NICUs causing neonatal infections are *Enterobacteriaceae* and *S. aureus.* Although many studies have emphasized transmission of MRSA (methicillin-resistant *S. aureus*), its susceptible variant MSSA is, depending on local epidemiology, generally a more frequent cause of neonatal infections (4,11,12). Neonatal bacterial infections are preceded by bacterial spread which involves both direct transmission by close interaction with visiting parents and HCWs and indirect transmission through medical (1-4). Proper hand hygiene or wearing disposable gloves and an apron have shown to be effective measures to reduce the incidence of infections in NICUs by 33-60% (5-7). To monitor the dynamics of potential pathogens in a NICU, carriage surveillance should be performed, as implemented at our NICU for *Enterobacteriaceae* and (both methicillin-susceptible and resistant) *S. aureus*. These advanced surveillance practices have become pivotal to the outbreak analysis performed in this study.

Outbreak analyses traditionally involve epidemiological analysis combined with microbial typing. Currently, traditional molecular methods for typing *S. aureus* include determination of the Multi-locus Sequence Type (MLST), the Variable Number Tandem Repeat (VNTR) signature of the staphylococcal protein A gene (*spa* typing), and the Multiple-Locus Variable number tandem repeat Analysis (MLVA)(8). Although MLVA has the largest discriminatory power (9), it covers only a fraction of the bacterial genetic material, and it is sensitive to influential sequencing errors.

With the advent of microbial whole-genome sequencing (WGS) information from the entire bacterial genome becomes available for typing and outbreak analyses (10). To facilitate such analyses, numerous new outbreak detection methods and applications have become accessible (11). Current examples of the application of WGS for outbreak analysis of *S. aureus* in neonates are mostly restricted to MRSA. Koser *et al*. first described WGS to confirm the epidemiological and phenotypic clustering of neonatal MRSA infections (12). Next, a suspected persistent outbreak of MRSA carriage among neonates was confirmed by WGS, possibly maintained by ongoing carriage in a HCW involved (13). In addition, WGS was demonstrated to enable differentiation between three contemporaneous MRSA outbreaks in a NICU (14), and also pointed towards international transmission of MRSA from NICUs (15). The sole example investigating MSSA, proposed that the application of WGS on neonatal infection isolates can also distinguish between transmission and exogenous introduction during an outbreak spanning multiple years (16).

Despite these appealing examples, uncertainty in WGS based outbreak analysis lies within the interpretation of sequence similarity, and the establishment of a threshold for transmission-level relatedness between bacterial isolates. Here, we performed WGS on MSSA isolates from both carriage surveillance and infections on a NICU, to more accurately reconstruct a period of presumed transmission events. To our knowledge, we are the first to deduce outbreak clusters being deviations from the background level of genomic diversity in the bacterial population as observed in surveillance of that particular clinical setting.

## Methods

### Setting and surveillance policies

This study was performed at a third level 18-bed neonatal intensive care unit (NICU) of the Radboud university medical center, Nijmegen, the Netherlands during 2014 and 2015. As part of bacterial surveillance purposes, once weekly a throat and rectal swab were taken from each neonate present at the NICU and cultured for 2 days on BD Columbia CNA agar with 5% sheep blood Improved II (Becton Dickinson) at 36°C. Colonies suspect for *S. aureus* that were positive in the slide coagulase test were tested for susceptibility to cefoxitin and mupirocin by agar diffusion. The first MSSA isolate cultured from a neonate was stored in glycerol 10% bouillon at −80°C for typing purposes. Surveillance culture results were discussed at a monthly meeting with representatives of the NICU and the department of Infection Prevention and Control to monitor colonization patterns.

### MSSA carriage and infection

MSSA carriage at a particular time point was defined as MSSA detected in at least one culture from a throat and rectal swab set. MSSA infection was defined as MSSA cultured from cerebrospinal fluid, blood, pleural fluid, wound swab, ocular swab or corneal scrape. Under certain conditions MSSA cultured from sputum was considered to represent possible or probable *S. aureus* infection (Supplementary methods 1). MSSA isolates cultured from infection sites were stored. Clinical characteristics of neonatal MSSA infections were manually extracted from corresponding patient records.

### Traditional outbreak investigations

During the study period, an outbreak was considered if closely related MSSA isolates were cultured from a throat and rectal swab set (carriage surveillance) or infection site in two or more patients within a month. Relatedness between isolates was assessed by agreement in traditional molecular *S. aureus* typing methods: by determination of the Variable Number Tandem Repeat signature of the *spa* gene (*spa* typing), and if *spa* type and temporal relationship raised suspicion for being a member of an outbreak, also by Multiple-Locus Variable number tandem repeat Analysis (MLVA) to determine the MLVA complex (MC) and underlying MLVA type (MT). If the time criterion was met, only isolates of equal MLVA types were considered to be sufficiently related to represent an outbreak. In retrospect, typing by whole-genome sequencing (WGS) was performed on a selection of the study isolates.

On occasion, when MSSA infections seemed to originate from carriage outbreaks, all involved health care workers (HCWs) were tested for MSSA carriage by a combined throat-nose swab and swabs from high frequency touched surfaces at the NICU were cultured. Those HCWs who carried outbreak *spa*/MLVA types were offered a five-day decolonization treatment and their corresponding isolates were subjected to WGS.

### Genomic analyses

The isolates selected for WGS differed per year. To estimate the level of genetic variation in background surveillance, for 2014 we included any typed MSSA isolate from neonatal carriage, plus the HCW carriage isolates of the outbreak MLVA types for WGS. For 2015, only the isolates with outbreak *spa* types were selected for WGS. A detailed description of the applied WGS methods is provided in Supplementary methods 2. In short, after DNA was extracted from cultured isolates by a cetrimonium bromide-based method, sequencing libraries were prepared for Illumina’s NextSeq500 platform using the NexteraXT kit (Illumina, San Diego, CA, USA). Reads were filtered and subsequently assembled into contigs by SPAdes (version 3.10.1) (17). Quality of the assembly was assessed by mapping the quality filtered reads back to the assembled contigs using bowtie2 (version 2.2.9) (18). Pairwise single nucleotide polymorphism (SNP) detection across all shared sequence regions between any pair of isolates was performed using kSNP (version 3.021) (19), followed by a custom script to obtain core high quality SNPs (hqSNPs) based on a specified sequence consistency and coverage. A phylogeny based on core hqSNPs was calculated using PhylML (20) and visualized using iTOL (version 4.1.1) (21). Minimum spanning trees (MSTs) were generated using python library networkx 1.11 and visualized using Cytoscape (version 3.5.1) (22). Because the Illumina NextSeq 500 platform yields paired-end reads of 150bp, the draft genomes were unfit to reliably span and verify the repeat-based typings that were previously assigned to the study isolates. The pairwise SNP analysis allowed us to valorize information from the “core” genomic regions shared by any subset of isolates from the cohort. In a subset of closely related isolates, the core genome is likely to be larger than in the total cohort, providing a substantially higher resolution for estimating relatedness. Because of this varying magnitude of the core genome, we measured relatedness between two isolates as the fraction of SNP differences across the shared genomic regions of a given subset that was present in the particular pair, denominated as SNP difference ratio. To account for possible technical SNP errors, introduced throughout the genomic analyses, we included several duplicates, including culture from frozen stocks as well as sequencing from libraries onwards.

### Study period from genomics perspective

Using the available genomic information, outbreaks during the study period were reconstructed. Outbreaks were now defined as groups of isolates with pairwise core hqSNP difference ratios between neighbors in the MST of the entire cohort below a particular threshold. This threshold could be no less than our technological SNP error rate (in the duplicates), yet was primarily based on the distribution of the number of pairwise core hqSNPs between MST-neighbors in the total cohort, including background surveillance. Through this approach, isolates were assigned to an outbreak cluster if they were genetically more similar than was expected based on the measured variation in background surveillance. While the threshold was selected arbitrarily, its appropriateness for the current cohort was subsequently assessed using epidemiological data. Epidemiological linkage was expected between the isolate pairs below the SNP threshold, while pairs above it ought to be unrelated.

## Results

### Surveillance

No methicillin-resistant *Staphylococcus aureus* (MRSA) was detected throughout the study period, which is expected with <1% carriage in the Dutch population (23).. Among neonates, in total 2296 surveillance swabs were collected, of which 313 (13.6%) were cultured positive for MSSA. Among the surveillance cultures 18.5% (214 out of 1154) of throat swabs were positive for MSSA, compared to only 8.7% (99 out of 1142) of rectal swabs. At a given surveillance time point, 39% of throat carriers were also positive in rectum culture, while 87% of rectal carriers were simultaneously positive in throat culture. The monthly number of throat swabs collected ranged from 74 to 122, while the proportion of positive MSSA cultures ranged from 2 to 41%. The majority of the 510 neonates involved in surveillance was screened only once before discharge from the NICU, and MSSA carriage was identified in 19.6% of them.

### Traditional outbreak analysis

Based on identification of equal MLVA types within one month time distance, five MSSA outbreaks were identified during the study period. The outbreak-MLVA types were observed simultaneously in both carriage as well as infection (Figure 1). The forty neonates who developed an infection with MSSA are further detailed in Supplementary table 1. On two occasions screening of surfaces and HCWs was performed, in May 2014 and November 2015. None of the cultured high frequency touched surface samples were positive for MSSA. Of the 377 screened HCWs 24% carried MSSA of whom 6 carried an outbreak MLVA type (Supplementary figure 1). The large variety of *spa* types identified in the entire cohort is displayed in Figure 2. We identified 68 unique *spa* types among 210 MSSA carriage isolates (121 first isolates from neonates and 89 from HCWs). The 93 isolates that underwent subsequent MLVA (49 neonates, 44 HCWs) showed 9 different MLVA clones, and 42 MLVA types. While most 2014 MSSA isolates were subjected to WGS, in 2015 isolates only outbreak *spa* types were selected for.

**Figure 1.**
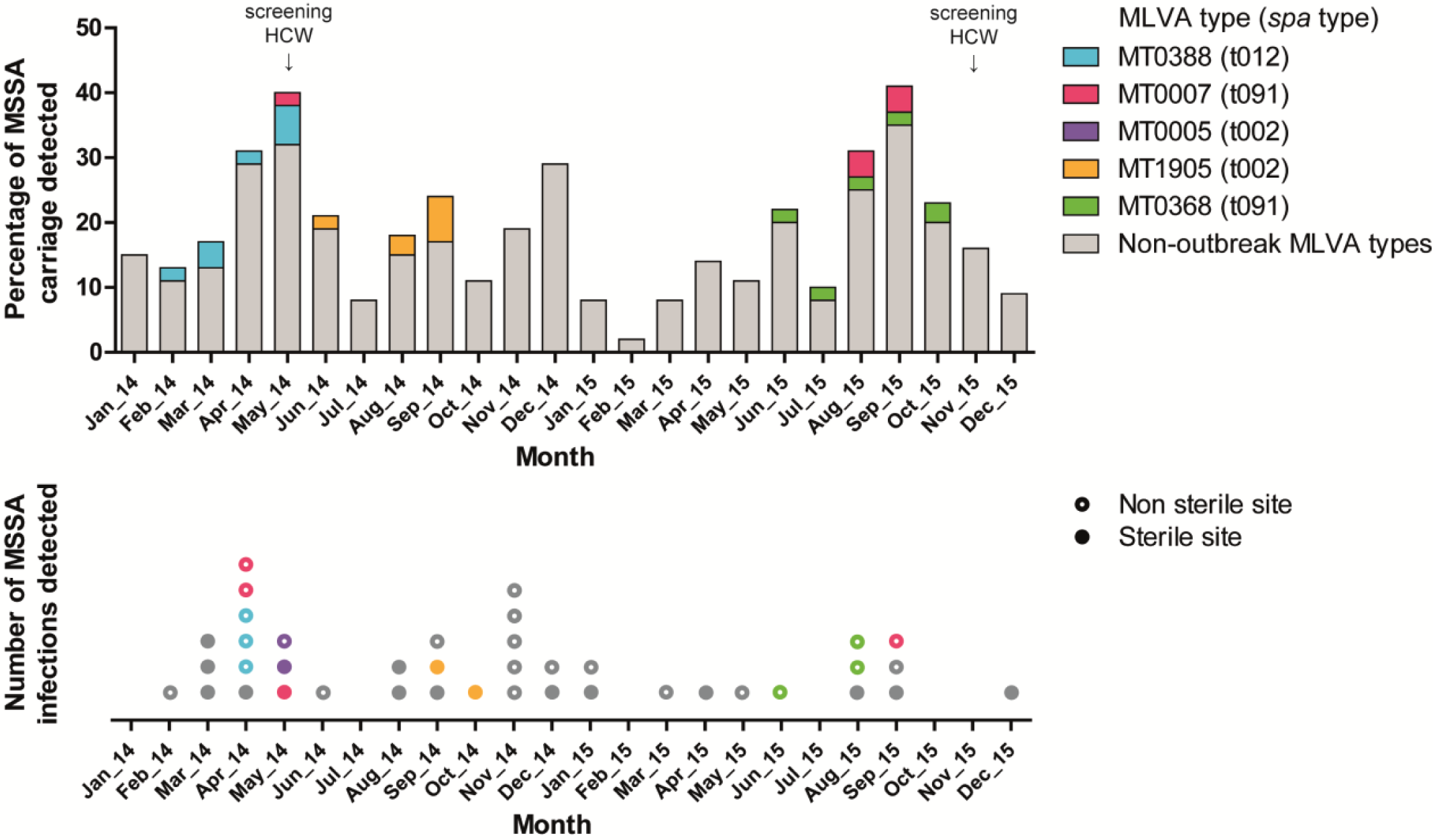
Traditional MSSA surveillance and outbreak view during the study period. The bars in the top panel indicate the monthly percentage of throat swabs positive for MSSA, collected in weekly neonatal carriage surveillance at the NICU over a 2 year period. The dots in the bottom panel indicate the absolute number of neonatal infections, according to specimen positive for MSSA. Members of the five presumed outbreaks are indicated by a given color. Vertical arrows indicate the two occasions on which cross-sectional screening of MSSA carriage in HCWs was performed. *Abbreviations*: MSSA: methicillin-susceptible *S. aureus*; MT: MLVA type; HCW: healthcare worker.

**Figure 2.**
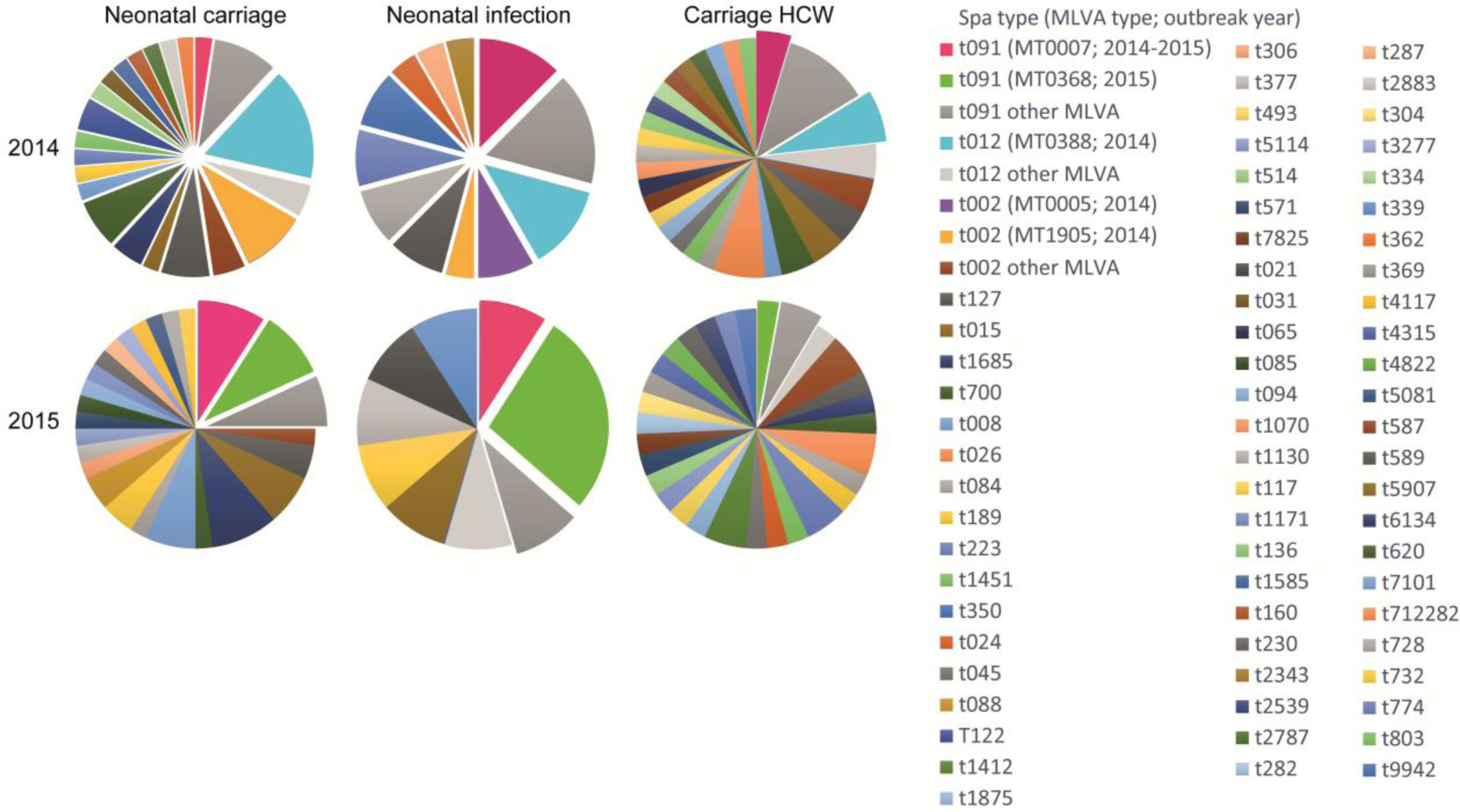
Population diversity and selection for whole-genome sequencing. Pie charts indicating the percentage of typed MSSA isolates assigned to a particular *spa* type or MLVA type, categorized by study year (rows) and origin (columns). Raised sections indicate the isolates that were eligible for whole-genome sequencing, which includes all neonatal isolates in 2014. *Abbreviations*: MSSA: methicillin-susceptible *S. aureus*; MT: MLVA type; HCW: healthcare worker.

### Genomic analysis

A total of 84 different MSSA isolates were sequenced, of which 78 from neonates (27 with MSSA infection) and 6 from HCWs. Six other eligible isolates were not stored and no longer available for WGS. The 84 selected WGS isolates represented 23 different *spa* types, and 58 of these isolates that had qualified for MLVA had been assigned to 5 different MLVA clones and 15 MLVA types. Quality thresholds for the genomic outbreak analyses were reached by 98% of the isolates. Sequenced isolates had a median sequence coverage of 88×, a median cumulative contig length of 2.75×10^6^ base pairs (bps) and a median paired-end read alignment to the contigs of 98%. Pairwise analysis of isolates identified 29,292 SNP-affected base positions present in the core genome of the total cohort, of which 8,575 (29%) were of high quality. The distance between the culture duplicates was 1 core hqSNP out of 8,575 hqSNP-affected base positions compared, and 0 core hqSNPs between the sequencing duplicates.

The concordance between *spa* typing, MLVA and genomic relatedness based on core hqSNPs in the total cohort is demonstrated in Figure 3. Stratification of isolates according to *spa* type (inner lane) was in perfect agreement with genomic relatedness. However, intense admixture of MLVA types (outer lane) is observed in the branch corresponding to *spa* type t091.

**Figure 3.**
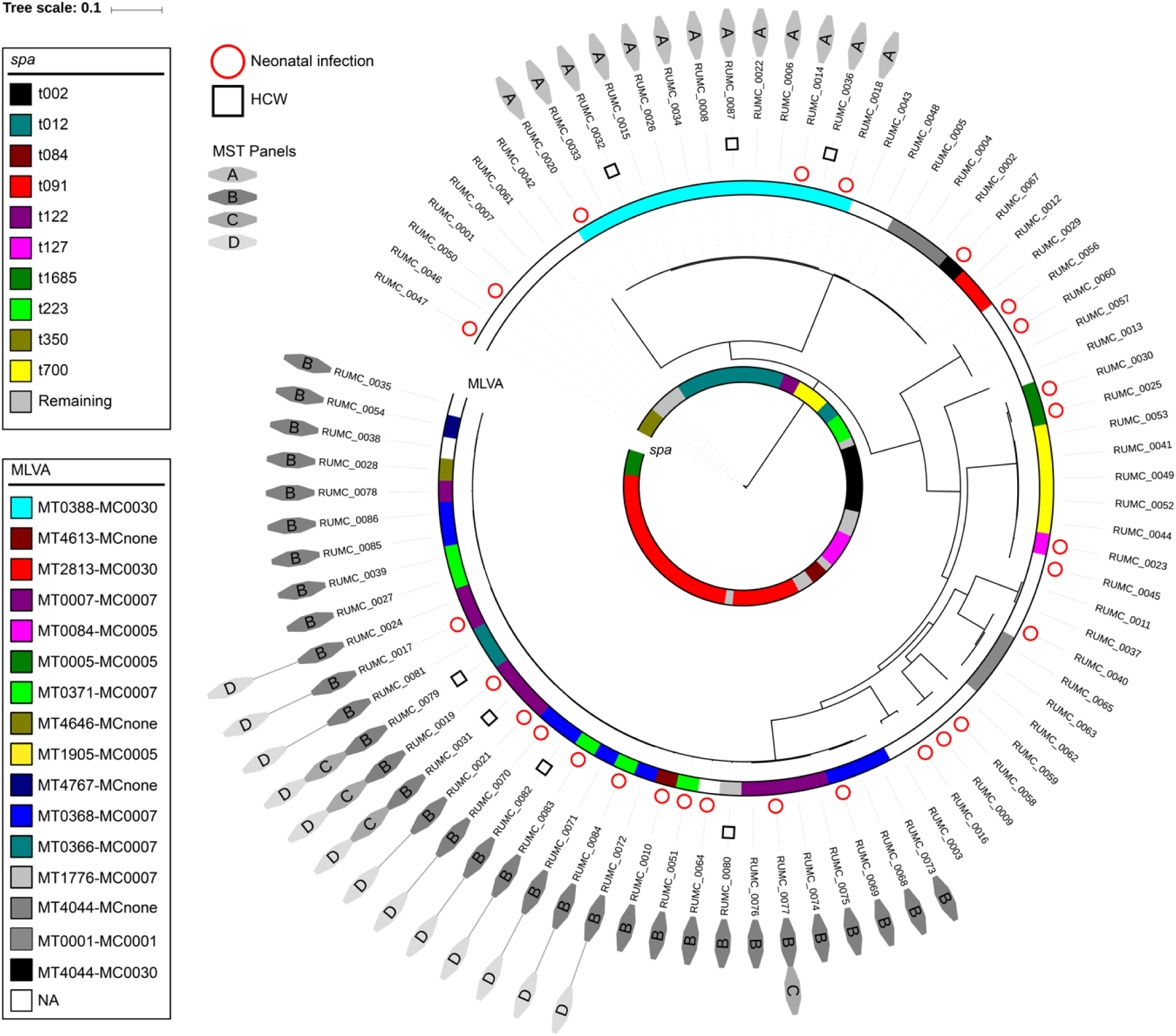
Concordance between traditional typing methods and WGS based phylogeny. The circular phylogenetic tree is based on 8,575 core hqSNPs among the sequenced isolates. Each branch represents an MSSA isolate. The colored lanes indicate *spa* type (inner lane) and MLVA type (outer lane). RUMC labels represent identifiers of deposited sequencing data. The labels A to D anticipate representation of the isolate in the MST panels of Figure 4. *Abbreviations*: MT: MLVA type; MC: MLVA clone; NA: not assigned; HCW: healthcare worker; MST: minimum spanning tree.

The extraction of subset specific core genomic regions significantly increased the resolution (the number of base positions screened for SNP differences) of our genomic outbreak analyses. For the 13 isolates of MLVA type 388 that were all regarded as members of a single outbreak by traditional analysis, a 6-fold increase in resolution by subset-analysis did not particularly change this perspective. Multiple HCWs were involved in this transmission chain (Figure 4A). However, a genomic close-up of the subset with disagreement between stratification according to MLVA type and genomic relatedness, displays SNP difference ratios of up to 0.32% between neighbors central to the MST (Figure 4B). Such large genomic differences make it highly unlikely that any of these MLVA types had been involved in long transmission chains. On top, it seems that the MLVA perspective has also induced erroneous separation of isolates that should have been attributed to a common outbreak. Examples are the extremely low SNP difference ratios between neighboring isolates of MT0371 and MT0368, or MT0007 and MT0366 in the MST. The re-assortment of isolates of these MLVA types by WGS clusters is supported by the common time periods in which they have been isolated. Figure 4C demonstrates how our subset-analysis alleviated concerns about a particular HCW. Contrary to what was concluded from MLVA, this HCW (isolate May_14 in Figure 4C) involved in a first outbreak (represented by isolate Apr_14) had reacquired a similar MSSA strain (isolate Oct_15), which was however unrelated to the second outbreak (represented by isolate Sep_15) with this MLVA type. Finally, knowledge of the genomic location of identified SNP differences between RUMC_0017 and RUMC_0071 (indicated by the arrow in Figure 4D) confirmed that these were not introduced by a single event involving concentrated footprints, but by multiple sparse events probably accumulated over time (Supplementary table 2).

**Figure 4.**
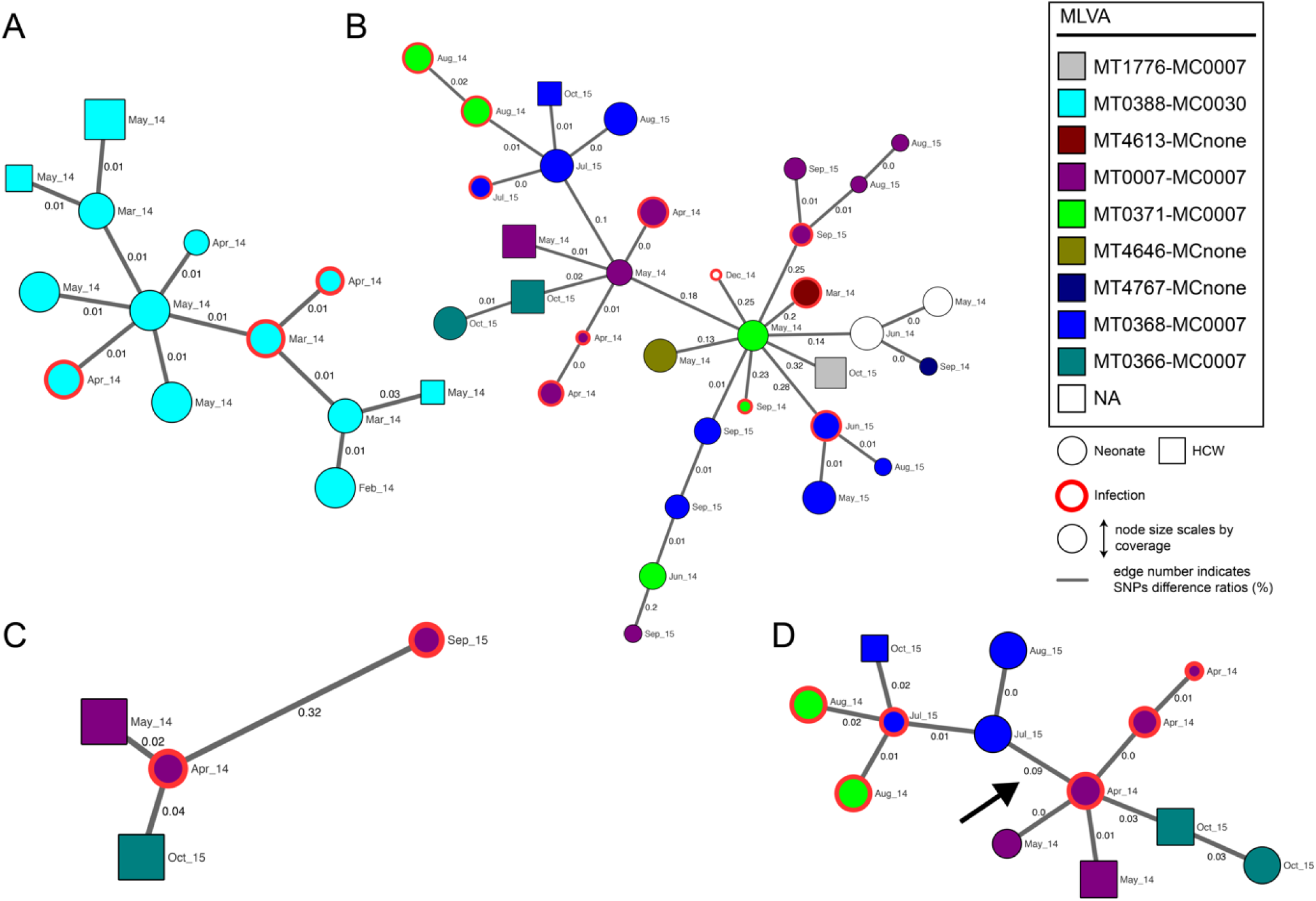
Minimum spanning trees of core hqSNP subset analyses. Nodes represent MSSA isolates which are labeled by month of isolation and colored according to traditional MLVA type assigned. Nodes of a subset are connected to their closest neighbors based on relative distances expressed in SNP difference ratios. Edges are not scaled. The arrow in panel D indicates the core SNP differences whose genomic locations are further specified. *Abbreviations*: hqSNP: high quality single nucleotide polymorphism; MT: MLVA type; MC: MLVA clone; NA: not assigned; HCW: healthcare worker.

### Study period from genomics perspective

The MSSA outbreaks were reconstructed using the available genomic information. Figure 5 demonstrates how, based on genomic distances in background surveillance, the threshold for transmission-level relatedness between neighbors in the MST was placed at a pairwise core hqSNP difference ratio of ≤ 0.03% (here equal to ≤ 3 hqSNP differences out of 8,575 SNP-affected base positions compared). Outbreaks were defined as groups of isolates that still met this criterion. To assess the appropriateness of this threshold for the current cohort we investigated the pairs of isolates around the cut-off for epidemiological linkage (Table 1).

**Table 1.**
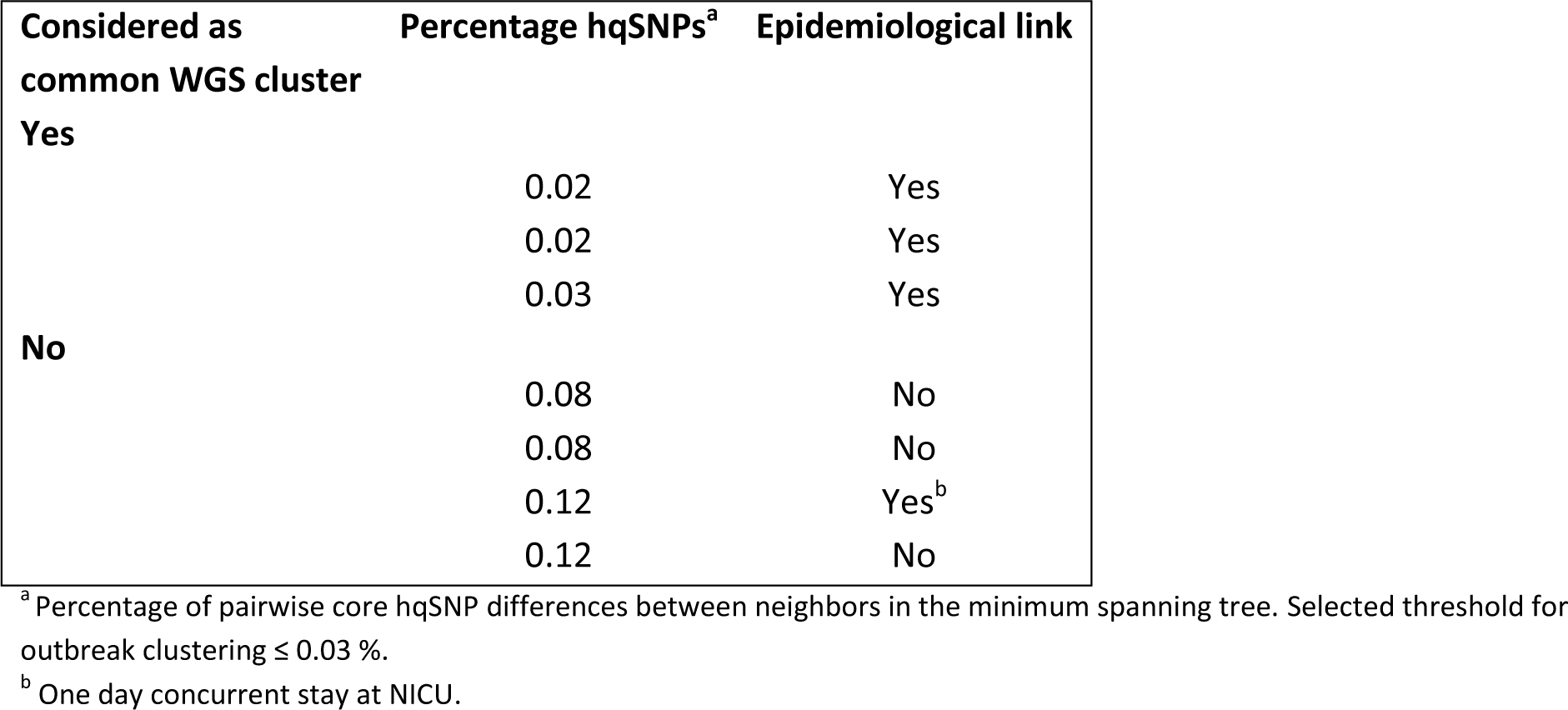
Epidemiological link between pairs of isolates around the WGS outbreak threshold.

**Figure 5.**
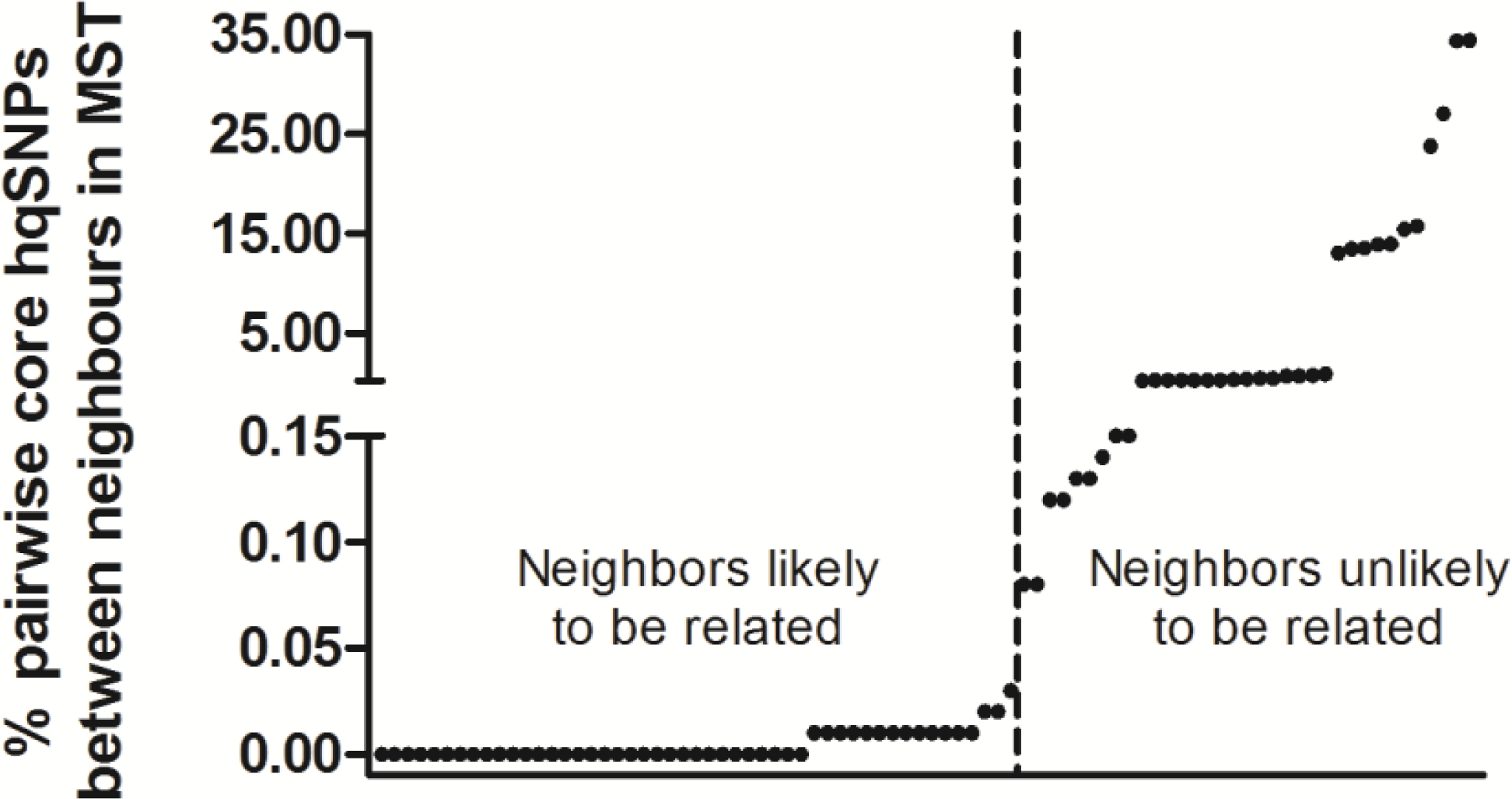
Threshold for transmission-level genomic relatedness. Dots indicate pairwise core hqSNP difference ratios between neighbors in the minimum spanning tree of the entire WGS cohort, ordered by increasing value. The threshold for transmission-level genomic relatedness was placed at ≤ 0.03% (y-axis), separating the MSSA neighbors into two groups as indicated by the vertical dashed line. *Abbreviations*: hqSNP: high quality single nucleotide polymorphism; MST: minimum spanning tree; WGS: whole-genome sequencing; MSSA: methicillin-susceptible *S. aureus*.

Figure 6 displays the outbreaks during the study period reconstructed according to our genomics approach. Compared to the traditional outbreak analysis (MLVA-typing plus time relation), the genomic analysis demonstrated ten more separate outbreaks, yet with shorter transmission chains, at least five more infections attributed to an outbreak (f.e. those in November 2014), and three infections attributed to a different outbreak (f.e. those in August 2015). The number of outbreaks identified increased from 4 to 10 in 2014 and from 2 to 4 in 2015, alongside an additional 2 outbreaks spanning both years (indicated in yellow and orange). On the other hand, the average chain length of outbreaks decreased from 5.5 to 3.9 neonates in 2014 and from 6.0 to 3.3 neonates in 2015. Both alterations support a better containment of transmission in 2015 compared to 2014 than was presumed by traditional outbreak analysis.

**Figure 6.**
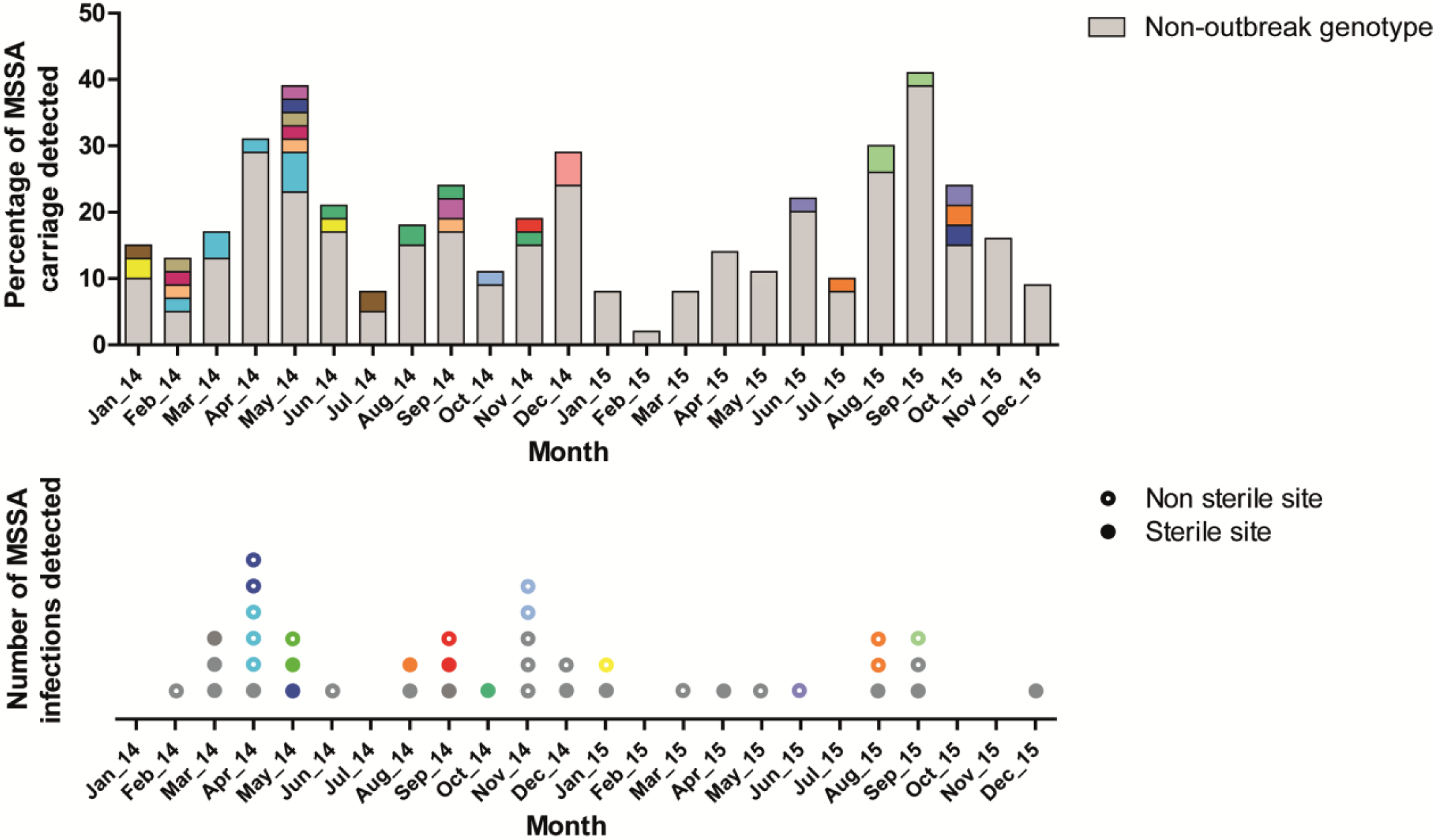
Neonatal MSSA outbreaks from a genomics perspective. Although panels display the same events as in Figure 1, members of outbreaks are now assigned according to our genomics approach. Colors are randomly assigned and not related to the previous figures. *Abbreviations*: MSSA: methicillin-susceptible *S. aureus*; MT: MLVA type.

## Discussion

Traditional outbreak tracing by *spa* typing and MLVA suggested that nosocomial transmission had contributed to neonatal MSSA infections in our patient cohort. Comparative genomics provided a large increase in resolution, especially in subset analyses, and yielded more accurate outbreak tracing than traditional typing methods. We based the genomic outbreak threshold on the background diversity in carriage surveillance, and acknowledged a substantially altered picture of outbreaks, which would have led to different outbreak measures.

One of the major issues that fueled this study, was the uncertainty about involvement of a particular HCW as a MSSA carrier in multiple outbreaks that included severe neonatal infections. Our genomic view alleviated these pressing concerns, and waived the working restrictions considered for this HCW.

A potential bias in our study is that, different from 2014, a selection of the MSSA isolates in 2015 was subjected to WGS, based on traditional typing methods. If all 2015 isolates had been sequenced, potentially more outbreak members would have been revealed, yet unlikely up to the same number as detected in 2014. Because we observed concurrent patterns of particular MSSA variants in carriage and infection, we believe that weekly neonatal surveillance and cross-sectional sampling of HCWs provided a good representation of the circulating MSSA population. However, one should keep in mind that such assessments are not designed to, and will not provide a complete capture of all transmission, and are likely to be affected by carriage density and chances of detection by screening technique used.

A relative limitation of our genomic method is that Illumina sequencing followed by *de novo* assembly does not allow for complete reconstruction of bacterial genomes, partly at the expense of repetitive elements. Although this issue can be overcome by long-read sequencing technologies such as Oxford Nanopore [10] or Pacific Biosciences single-molecule real-time sequencing [10, 30], it is unsure whether genetic information from such complex regions would improve outbreak tracing. Although the genomic analyses as performed in this initial study are computationally demanding, they served well for sorting unresolved transmission chains. Especially if the surveillance-based concept presented in this study would be proven applicable to other outbreak settings as well, it would be worthwhile to further automate the genomic analyses, including outbreak threshold selection and outbreak subset analysis.

The strength and novelty of the current study lies in the combination of surveillance practices and particular genomics methods. Although surveillance of MSSA carriage is no common practice, its initiation at our NICU quickly paid off when a rise in MSSA infections was observed and could partly be attributed to nosocomial transmission. In addition, the availability of isolates from the circulating MSSA carriage population provided us the unique opportunity to investigate whether knowledge of background genetic diversity could tackle the issue of determining thresholds for genomic outbreak calling. Two aspects of our genomic methods were designed to maximize the proportion of the genomes that could be utilized for comparison. First, because genomes were reconstructed *de novo* from the sequencing reads, identification of genetic regions was not restricted to those present on a reference genome. Second, in addition to the SNP analysis on the core genome of the entire cohort, the applied k-mer based pairwise SNP analysis allowed us to substantially expand the genomic regions for comparative subset analysis. Compared to recently applied core genome and whole genome MLST methods that rely on alleles (24, 25), our method is more dynamic as it includes any genomic region in common between a pair of isolates, including non-coding regions. This “zooming in” subset analyses appeared not only advantageous because of an absolute increase in base positions compared, but there was also more discriminatory signal in the accessory regions. We observed that for the same pair of isolates, an expansion of the core genome generally increased the percentage of hqSNPs detected, probably because genomic regions that are not universally essential to *S. aureus* survival allow for more variability. With this increased resolution the neighbors in the MST stayed put, yet their degree of relatedness became better specified. Finally, to account for the possibility that multiple SNP differences were in fact introduced by a single event like horizontal gene transfer, we designed a method to determine the distribution of SNP locations across multiple bacterial genomes without using a reference genome.

For *S. aureus* an MLVA type is determined by the pattern of 24bp repeats within eight genetic regions (V09_01-V61_01-V61_02-V67_01-V21_01-V24_01-V63_01-V81_01). An explanation for the excessive admixture of four MLVA types compared to the WGS-based stratification in part of our MSSA population could be that these MLVA types were defined by highly similar repeat patterns (MT0368: 14-1-1-5-1-11-8-6; MT0007: 16-1-1-5-1-11-8-6; MT0366: 18-1-1-5-1-11-8-6; MT0371: 16-1-1-5-1-11-7-6). This suggests that either the MLVA typing method is susceptible to errors, or that MLVA repeat regions are simply not always representative for the remainder of the genome. The knowledge of epidemiological links between isolates of our study cohort made us confident that the genomic outbreak view was more plausible and therefore more reliable than MLVA type clustering. While in line with previous reports *Spa* typing concurred with WGS-based phylogeny (26, 27), it lacked the required resolution as exemplified by the five separate outbreaks that were identified under *spa* type t091 using comparative genomics.

Because our specified genomic methods including particular SNP confidence requirements have not been applied before, the selected SNP threshold cannot be directly compared to that in other genomic outbreak studies. However, previous WGS outbreak studies on *S. aureus* infections among neonates accepted isolates to belong to an outbreak cluster if they differed dozens of core genomic sites, compared to our restriction at 3 hqSNPs. The admixture of isolates from multiple *spa* types (28) did not occur in our study. In most studies, a relatively devoid group of isolates was arbitrarily considered as separate outbreak (12-16). Sometimes WGS analysis resulted in a further separation, but never in re-assortment of outbreak isolates like we observed. While several studies referred to clonal mutation rates reported in unrelated cohorts to infer transmission-level relatedness, Coll *et al*. based their outbreak threshold of ≤50 core SNP differences in a monophyletic group of isolates on those observed within multiple isolates from single patients, and the maximum number of SNP differences between isolates within the largest epidemiologically trusted outbreak in their cohort (29). Although accuracy and resolution may further benefit from knowledge on the local circulating carriage population, WGS benchmark initiatives could provide an appropriate platform for comparing the effects from different genomic outbreak methodologies on the outbreak assessment for a particular cohort (30). While our threshold for genomic outbreak calling was supported by the epidemiological linkage between isolates, this threshold was put rather low, demanding tight genomic relatedness to meet clustering. These settings resulted in a genomic view of more and shorter outbreaks compared to traditional typing methods. Although we cannot exclude that the selected threshold was too strict, the re-assortment of outbreaks did not lead to indistinct singletons but rather separations into more homogeneous sub-clusters, still suggestive of a credible genomic outbreak reconstruction.

The genomic view revealed pairs of isolates that were collected months apart but displayed identical core genomes. Given the vast genetic diversity observed in the circulating MSSA population, we considered that these isolates were very likely to be closely related. Likewise, previous studies identified many unsuspected transmission clusters upon re-assessment of *S. aureus* cohorts using WGS (28, 29). Accordingly, in our genomic view time-linkage was not regarded as a prerequisite for outbreak clustering. This contributed to the increased number of outbreaks detected and the larger number of infections being attributed to nosocomial transmission.

Despite the non-exhaustive typing strategy applied in this study, MSSA isolates from 19 out of 40 neonatal MSSA infections were demonstrated likely to be members of transmission chains involving other neonates. Although prevention of nosocomial transmission of micro-organisms does not necessarily ban all neonatal infections, the attention for MSSA transmission seemed to have resulted in less spread and infections in the second year of the study. By our genomic view, infection prevention measures sorted an even larger reduction of outbreaks than presumed by traditional outbreak analysis. In addition, the outbreaks in 2015 appeared more modest, which could have waived the incentive to perform a second screening of MSSA carriage among HCWs.

Future studies are required to assess whether the current approach can again be successfully applied in similar as well as different outbreak settings involving other pathogens. Particularly because the rate at which mutations accumulate varies between bacterial species and lineages (31). In line, within our cohort we noticed that the degree of SNP variation seemed to differ between lineages (Supplementary figure 2). It is unsure at which level such variation would violate the representativeness of a single outbreak threshold for an entire cohort. A related question is how extensive (in terms of density, but also proximity in place and time) background surveillance needs to be in order to identify an adequate threshold for transmission-level genomic relatedness.

### Conclusions

In this study, WGS-based reconstruction of MSSA outbreaks was clearly superior to that based on traditional typing methods. As accurate insight in transmission is crucial for proper selection and evaluation of interventions for containment, we recommend WGS as the preferred method for assessment of bacterial outbreaks. Consensus needs to be reached on what would be needed for real-time genomic surveillance to become the leading method for identification of nosocomial outbreaks (32, 33).

## Declarations

### Ethics approval and consent to participate

Not applicable.

### Consent for publication

Not applicable.

### Availability of data and materials

The dataset supporting the conclusions of this article is available in the Sequence Read Archive repository; BioProject accession number PRJNA523569; https://www.ncbi.nlm.nih.gov/sra/PRJNA523569.

### Authors’ contributions

AC and JC wrote the first version of the manuscript. AC and JC performed the data collection. JZ performed the laboratory assays. AC performed the epidemiological data analyses and JC performed the bioinformatics analyses. HW and JH initiated and supervised the study. All authors read, contributed to, and approved the final manuscript.

## Acknowledgements

Not applicable.

## Supplementary Methods

### Supplementary Methods 1. Definition of MSSA infection

In case a neonate was suspected for a respiratory infection, a standard work-up including blood cultures was performed. If sputum was the most invasive specimen in which *S. aureus* had been identified, we retrospectively stratified the likelihood of a true *S. aureus* infection to be unlikely, possible, or probable. *S. aureus* infection was denominated unlikely if a respiratory *S. aureus* infection was not adequately covered by a subsequently administered antibiotic treatment regimen, if a more evident infectious cause was identified, if clinical signs at specimen collection were deemed unrelated to infection based upon the subsequent clinical course, or if the culture result was explicitly considered as not related to infection by the treating physician. Unlikely cases were removed from further analyses of *S. aureus* infections. Among the remaining sputum positive cases *S. aureus* infection was denominated probable if the neonate was in respiratory distress, and had a blood CRP-level >20mg/L or a lung infiltrate on chest X-ray. The remaining cases were classified as possible *S. aureus* infection.

### Supplementary Methods 2. WGS methods

#### DNA isolation and library preparation

Selected MSSA isolates were cultured overnight at 36°C on BD trypticase soy agar II with 5% sheep blood. Bacterial DNA was extracted by a CTAB-based method. Colonies were resuspended in 400 µL TE buffer (10 mM Tris pH8.0, 1 mM EDTA) and 50 µL of 10 mg/mL lysozyme was added. Samples were incubated for 60 min at 37°C, whereafter 75 µL of 0.7 mg/mL proteinase K in 10% SDS was added followed by an incubation for 10 min at 65°C. The samples were mixed with 100 µL of 5M NaCl and 100 µL CTAB/NaCl solution (1% N-cetyl-N,N,N,-trimethyl ammonium bromide in 0.7 M NaCl) and incubated for 10 min at 65°C. DNA was further isolated using chloroform/isoamylalcohol extraction. DNA was precipitated from the aqueous phase by adding an equal amount of 2-propanol and subsequent incubation for 20 min at −20°C. Samples were centrifuged for 10 min at 11.000*g*. DNA pellets were washed with 1 mL cold 75% ethanol and centrifuged for 5 min at 11.000*g*. DNA pellets were air-dried for 15 min at room temperature and dissolved in 100 µL TE buffer. DNA samples were quantified using the QuantiFluor dsDNA system (Promega, Madison, WI, USA). A fragmented genomic DNA library was prepared using a NexteraXT DNA sample preparation kit (Illumina, San Diego, CA, USA). Subsequent sequencing was conducted in a paired-end 2 × 150bp mode using an Illumina NextSeq500 sequencer (Illumina, San Diego, CA, USA).

#### Read trimming and assembly

Sequence reads containing Nextera XT adapters and low quality regions were detected and removed by Trim_galore (version 0.4.1) (1), using *“--length 150 --stringency 12”* settings. Reads with tailing artefacts (stretches of minimally 6 A’s or 6 G’s introduced by sequencing chemistry) were removed using a custom python script. Reads with lengths of either 150 or 151 were used for de novo assembly. Prior to assembly the read coverage was estimated by dividing the number of nucleotides on the filtered reads by the length of *Staphylococcus aureus* strain RIVM1295 (RefSeq:NZ_CP013616.1). Reads were randomly subsampled to 120x coverage if coverage exceeded. To obtain high quality assemblies, isolates with an average coverage < 20x were discarded from further analyses. Reads were assembled into contigs with SPAdes 3.10.1 (2) at default settings and using k-mer sizes 21, 41, 61, 81 and 101. Contigs were annotated using PROKKA (3). Read depth at each position was determined by mapping reads to the contigs using Bowtie2 (version 2.2.9) (4) *“--no-unal –no-discordant”*, followed by Samtools (version 1.3.1) (5) depth estimation. The Percentage of reads mapped to the contigs is used as measure of assembly quality. Prior to outbreak analyses contigs with a mean coverage depth < 20x and/or contigs with length < 500 bp were discarded.

#### SNP handling

Single nucleotide polymorphisms (SNPs) were determined by k-mer extraction from qualified contigs, using kSNP (version 3.021) (6). The optimal k-mer size for *S. aureus* of 19bp was determined using Kchooser (6). High quality SNPs (hqSNPs) were obtained by mapping the k-mers to the contigs using Bowtie2 (4) “--all”. k-mers that map perfectly multiple times and/or not map perfectly were excluded for further analyses. A custom python code derived each k-mer location from the bam files and determines the exact SNP position on the contigs. Prokka annotation file in gff format is parsed to match the SNP location, adds annotation to be able to determine if SNP location is on coding or non-coding region of the genome. Previously determined read depth is used to filter SNPs with ≥ 20x read depth, resulting in hqSNPs. Core hqSNPs were defined as hqSNP positions located at genomic regions that were present in all strains selected for a particular analysis. Phylogeny by a maximum likelihood tree was inferred from all concatenated core hqSNPs per strain. The model was determined by using jModelTest 2.1.10 (7) resulting by applying PhylML (8) with custom best model “-d nt -n 1 -b 0 -m 012314 -f m -c 1 -- no_memory_check -o tlr -s BEST”. Visualization of the phylogenetic tree was done using iTOL v4.1.1 (9). Kruskal’s Minimum Spanning Trees (MST) (10) were calculated from pairwise core hqSNP differences between all strains, using networkx 1.11 in Python 2.7.5, and visualized using Cytoscape (3.5.1) (11).

## Supplementary Tables

**Supplementary Table 1.**
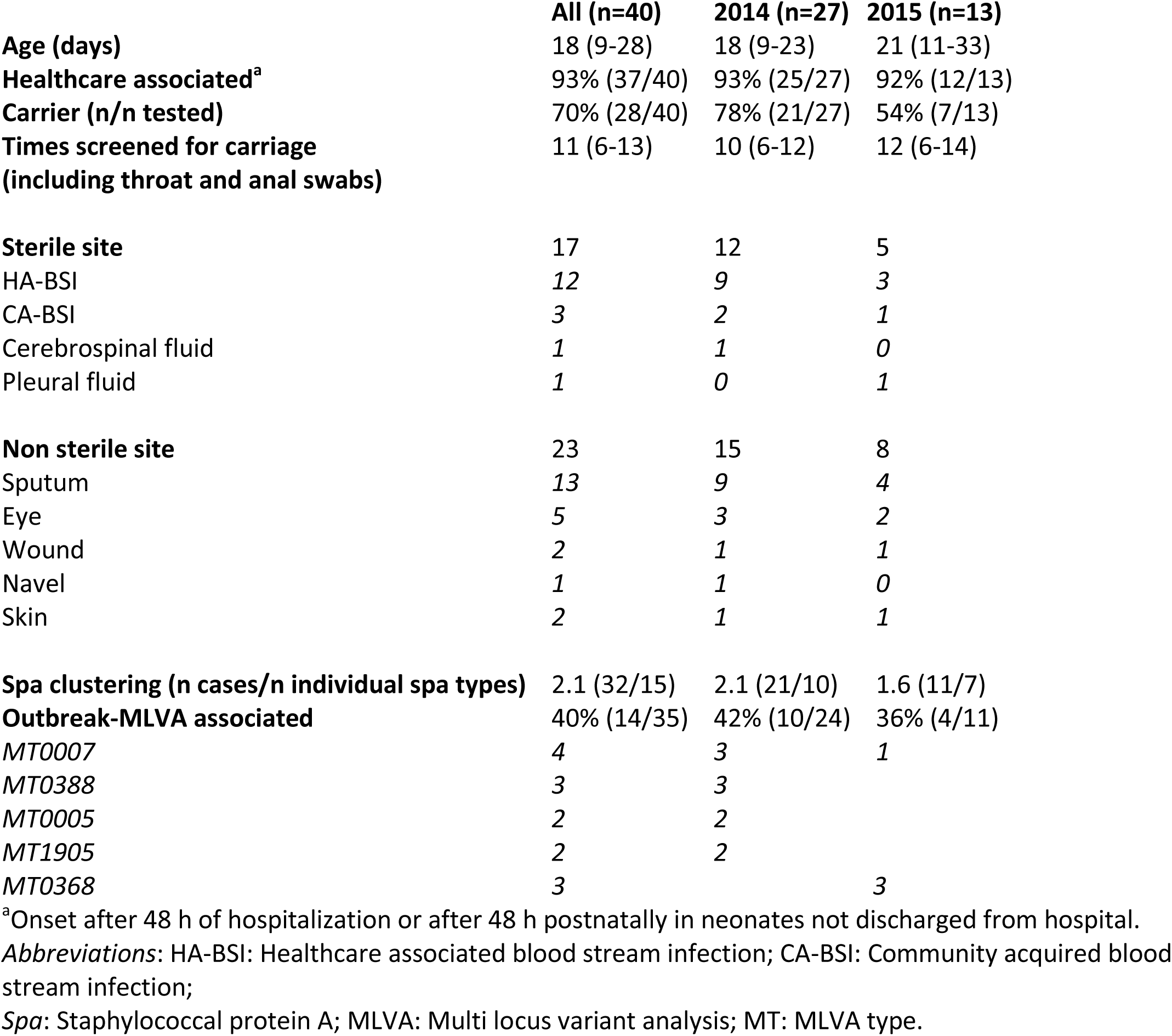
Neonatal MSSA infections.

**Supplementary Table 2.**
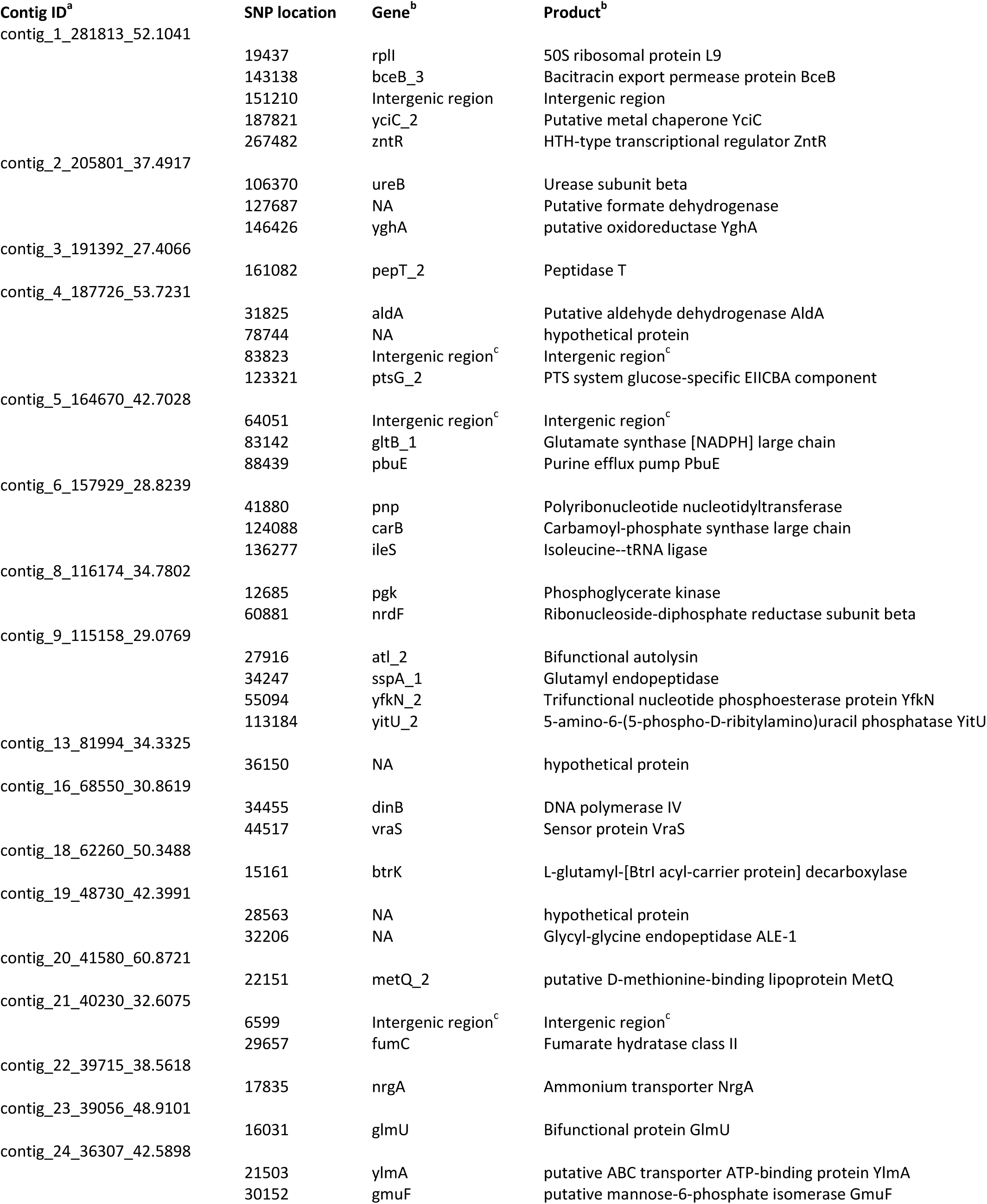
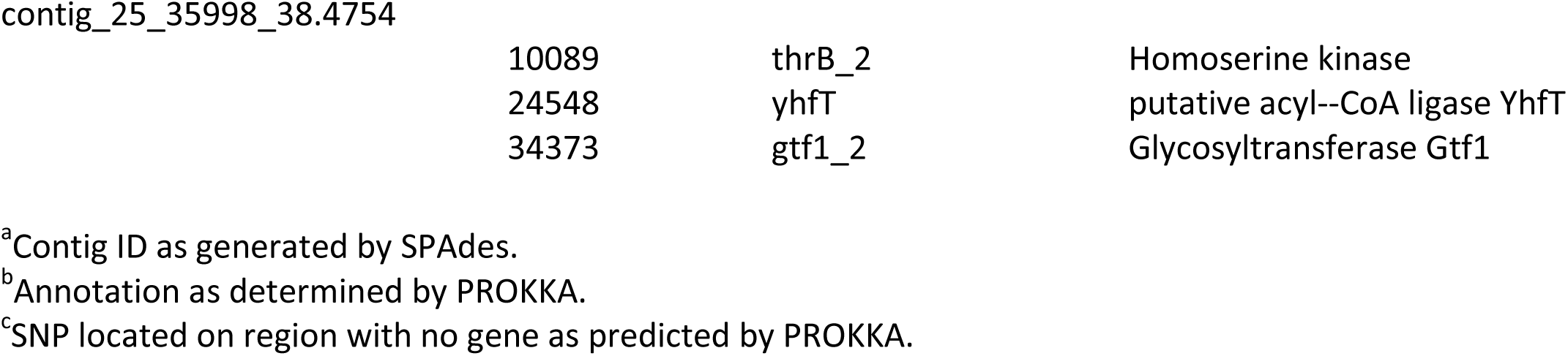
hqSNP locations between isolates RUMC_0017 and RUMC_0071.

## Supplementary Figures

**Supplementary Figure 1.**
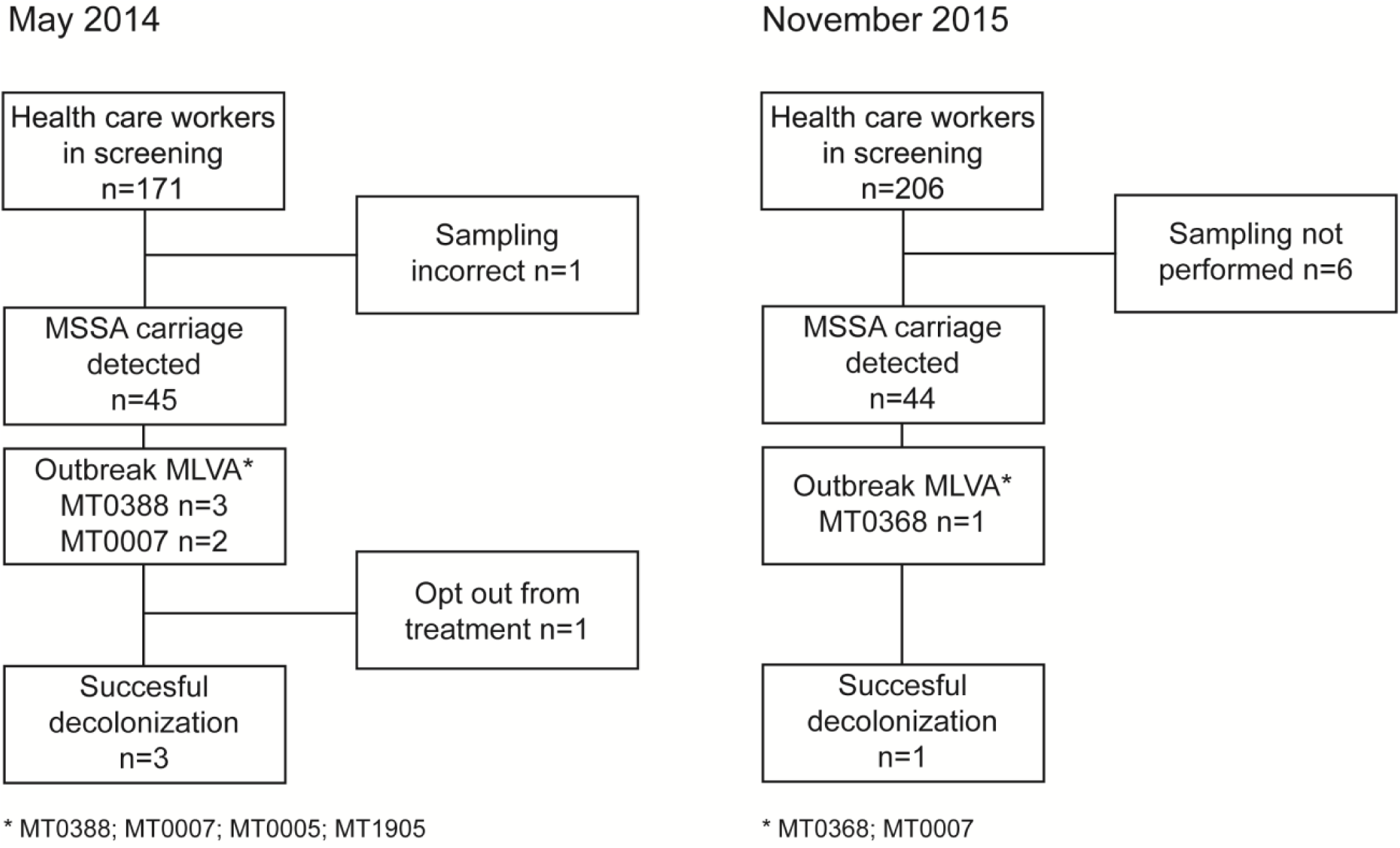
Flow chart screening HCWs.

**Supplementary Figure 2.**
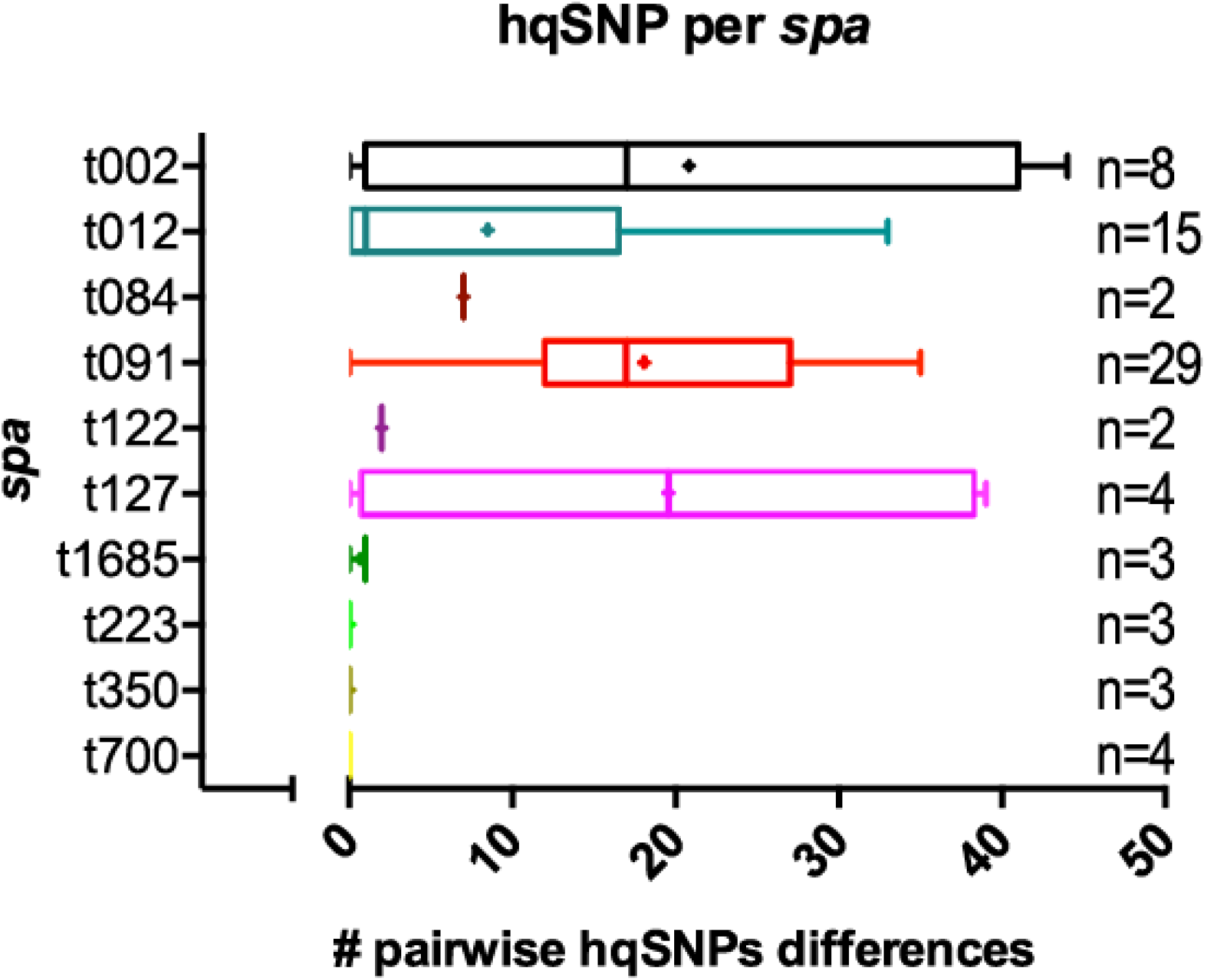
Diversity in percentages of pairwise core hqSNPs between members of particular *spa* types.

